## Supplementary figures and images for "A stochastic world model on gravity for stability inference"

### Figure1.png

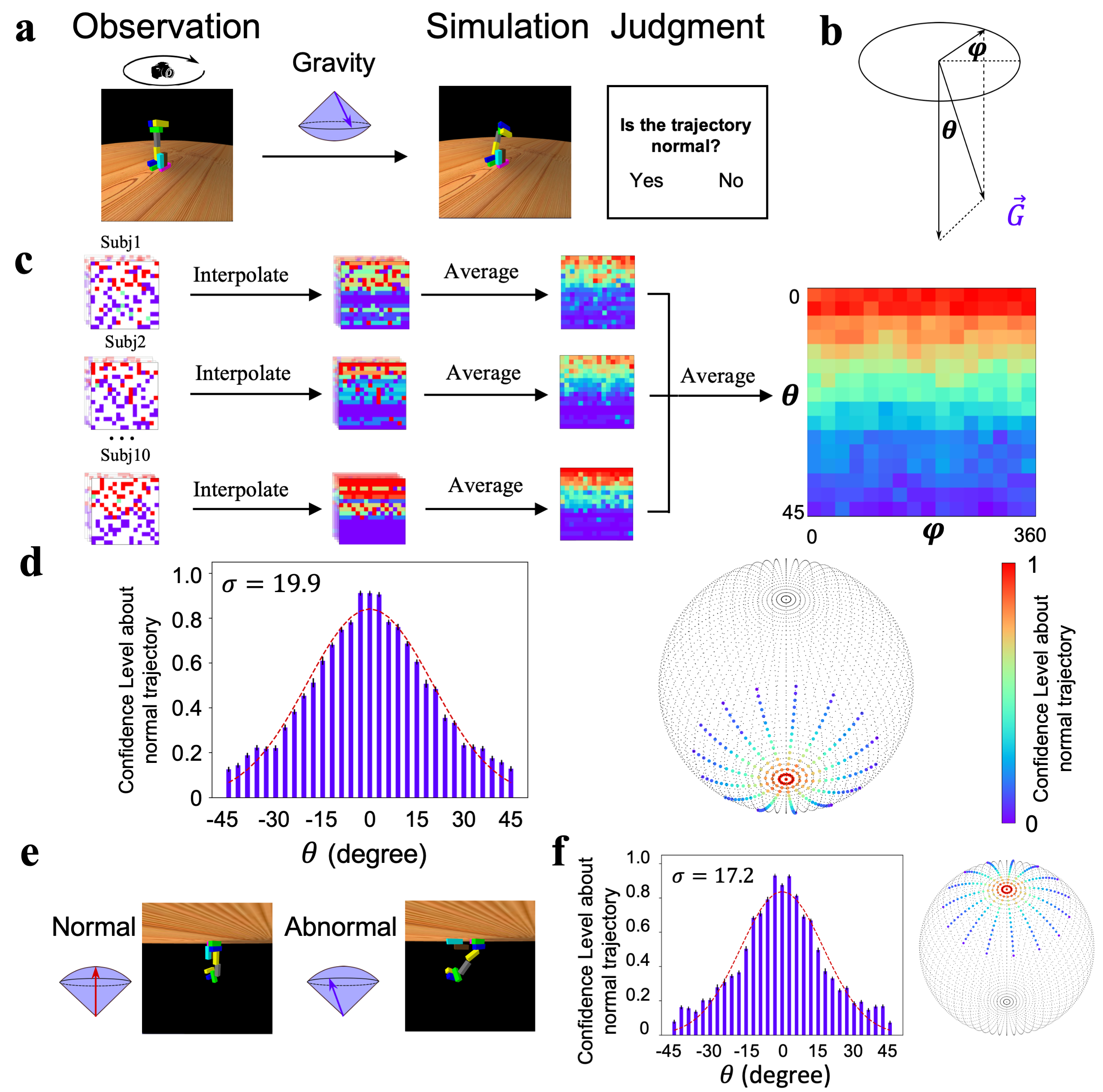

### Figure2.png

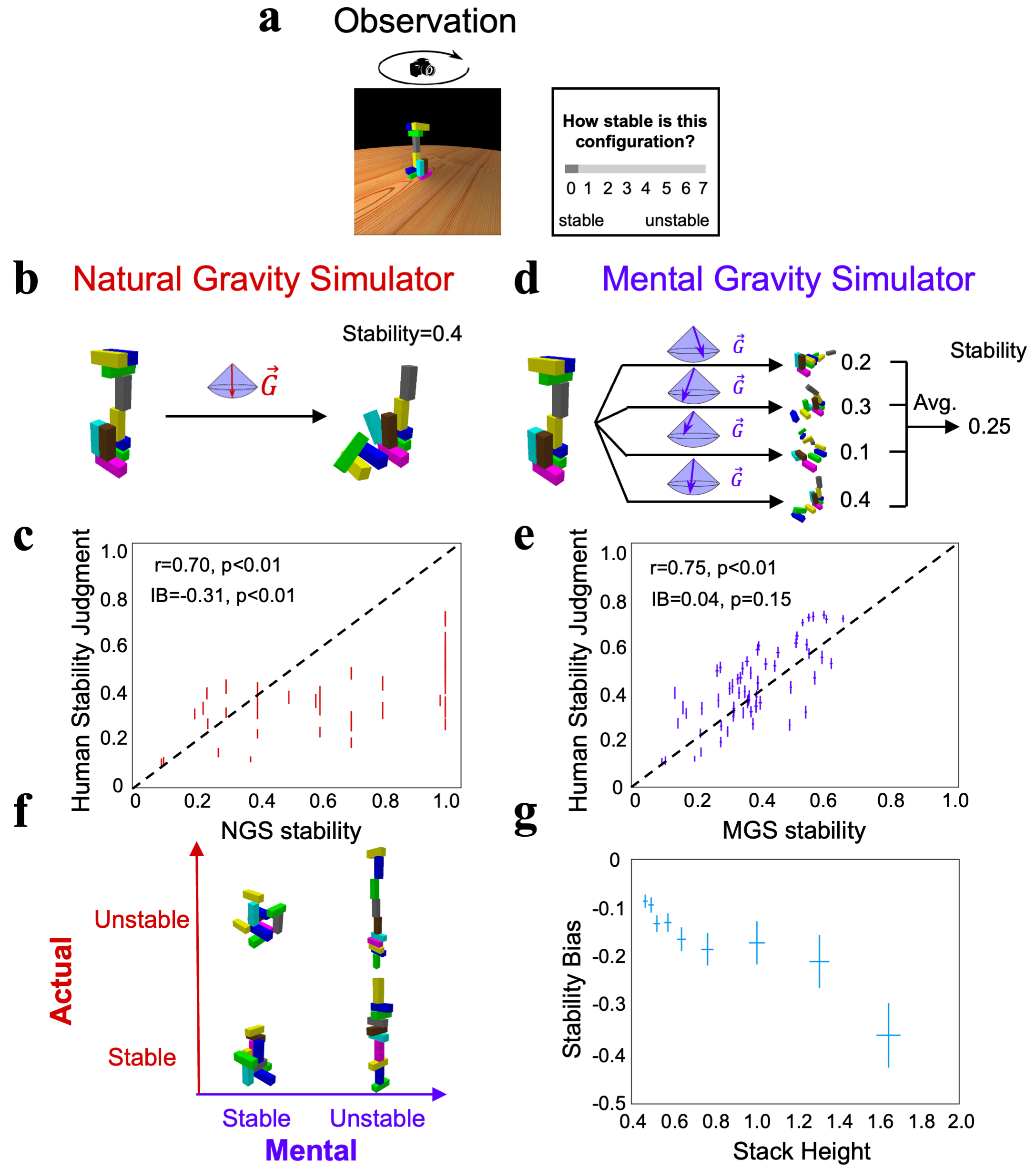

### Figure3.png

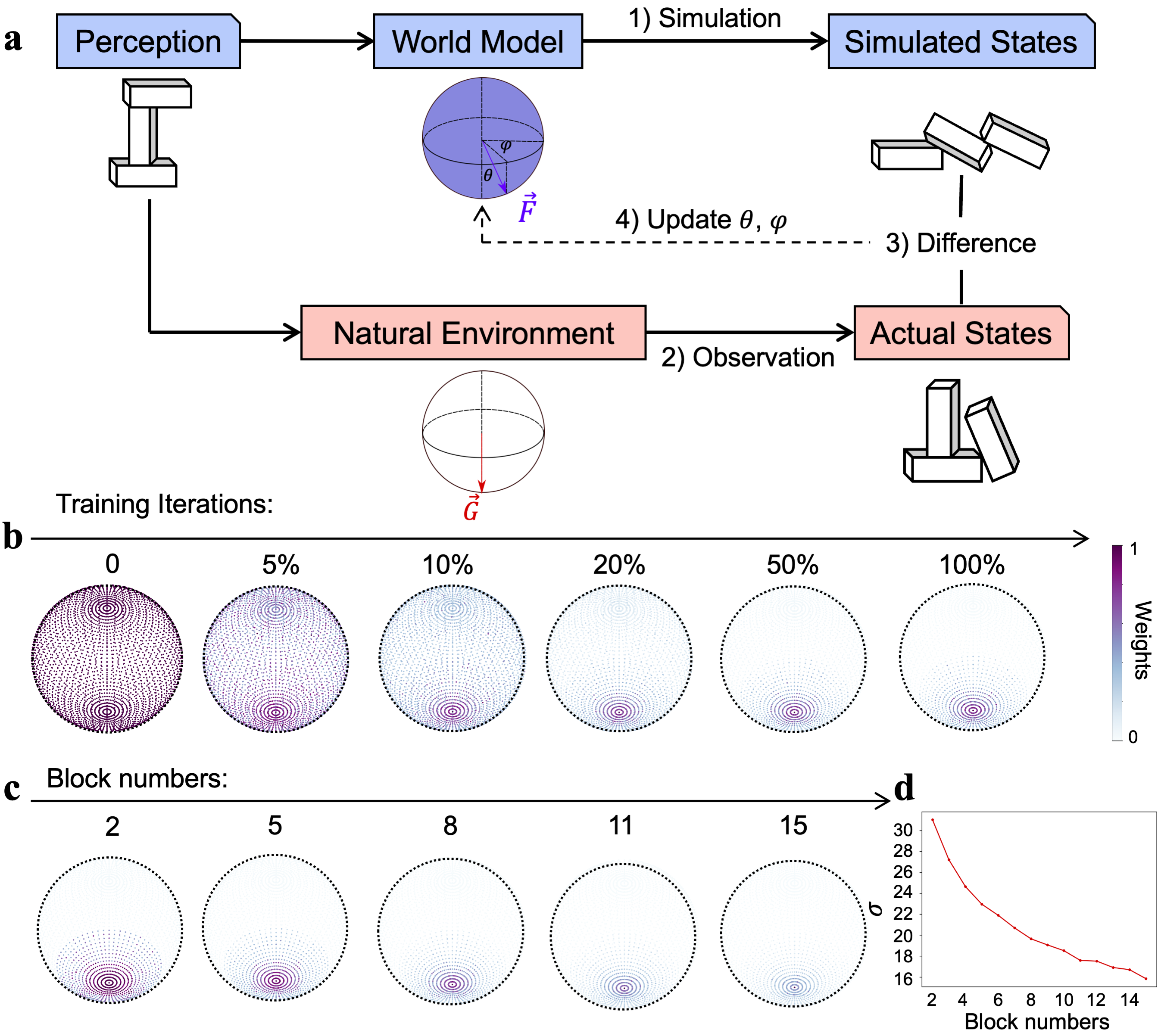

### Figure4.png

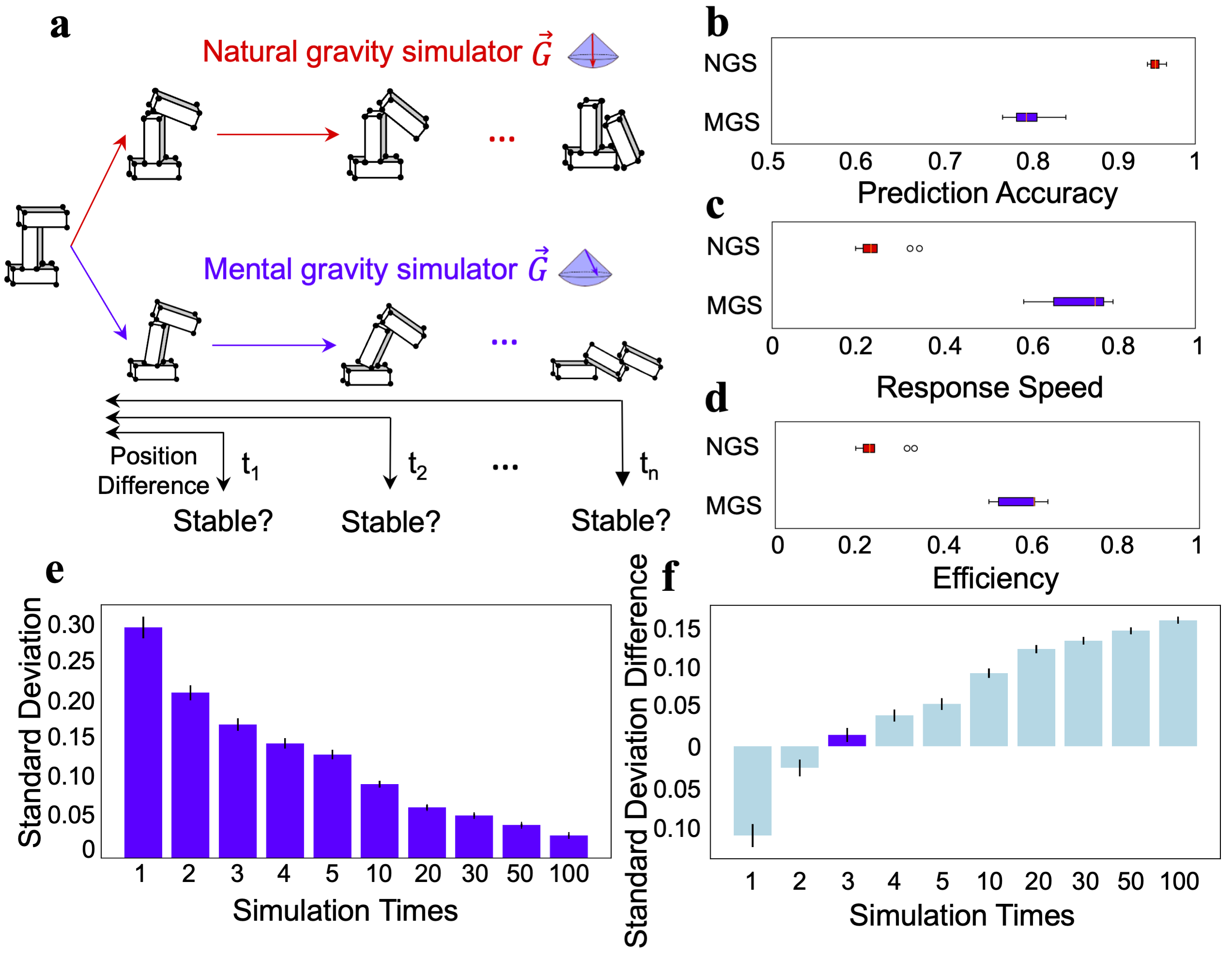

### supFigure1.png

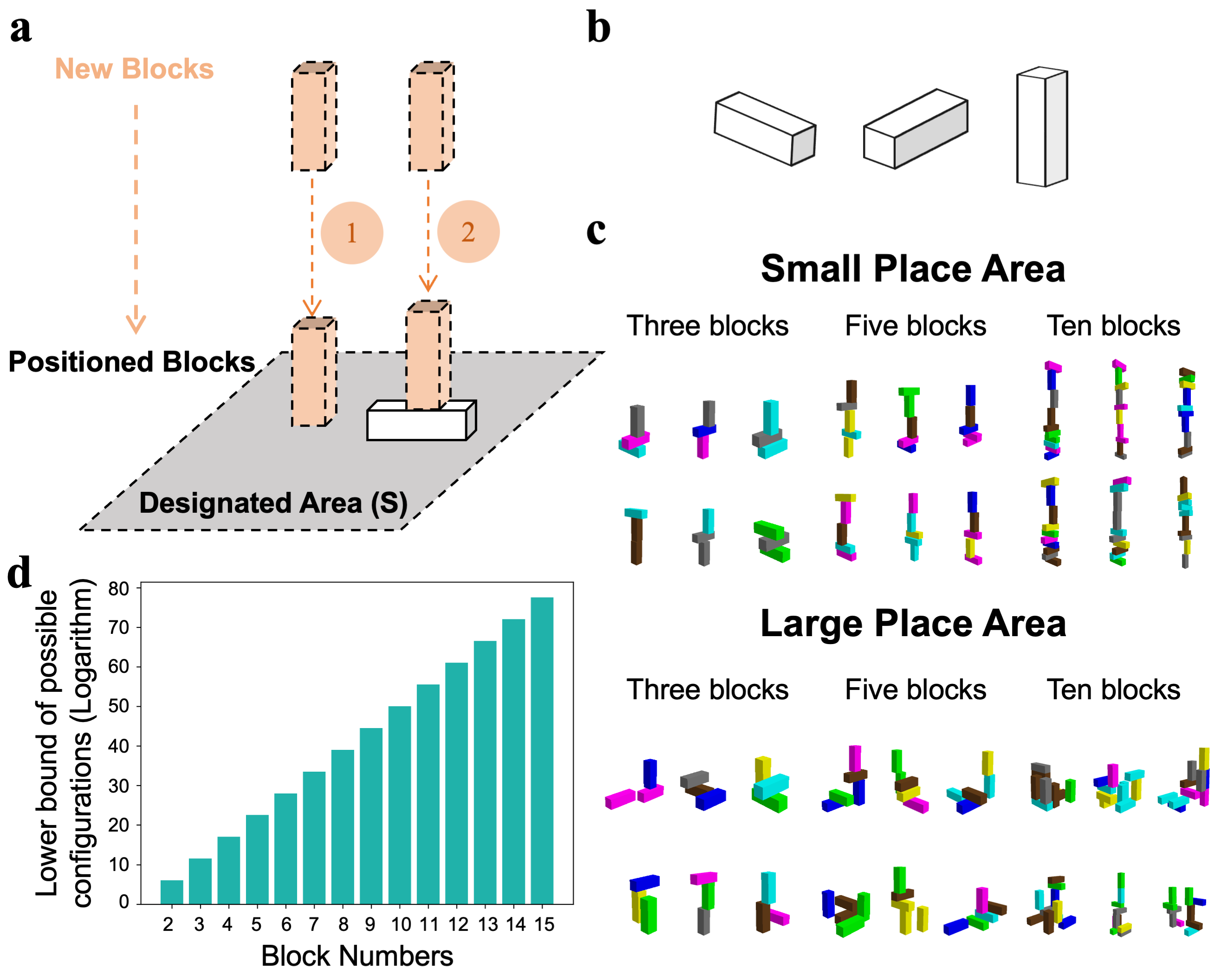

### supFigure2.png

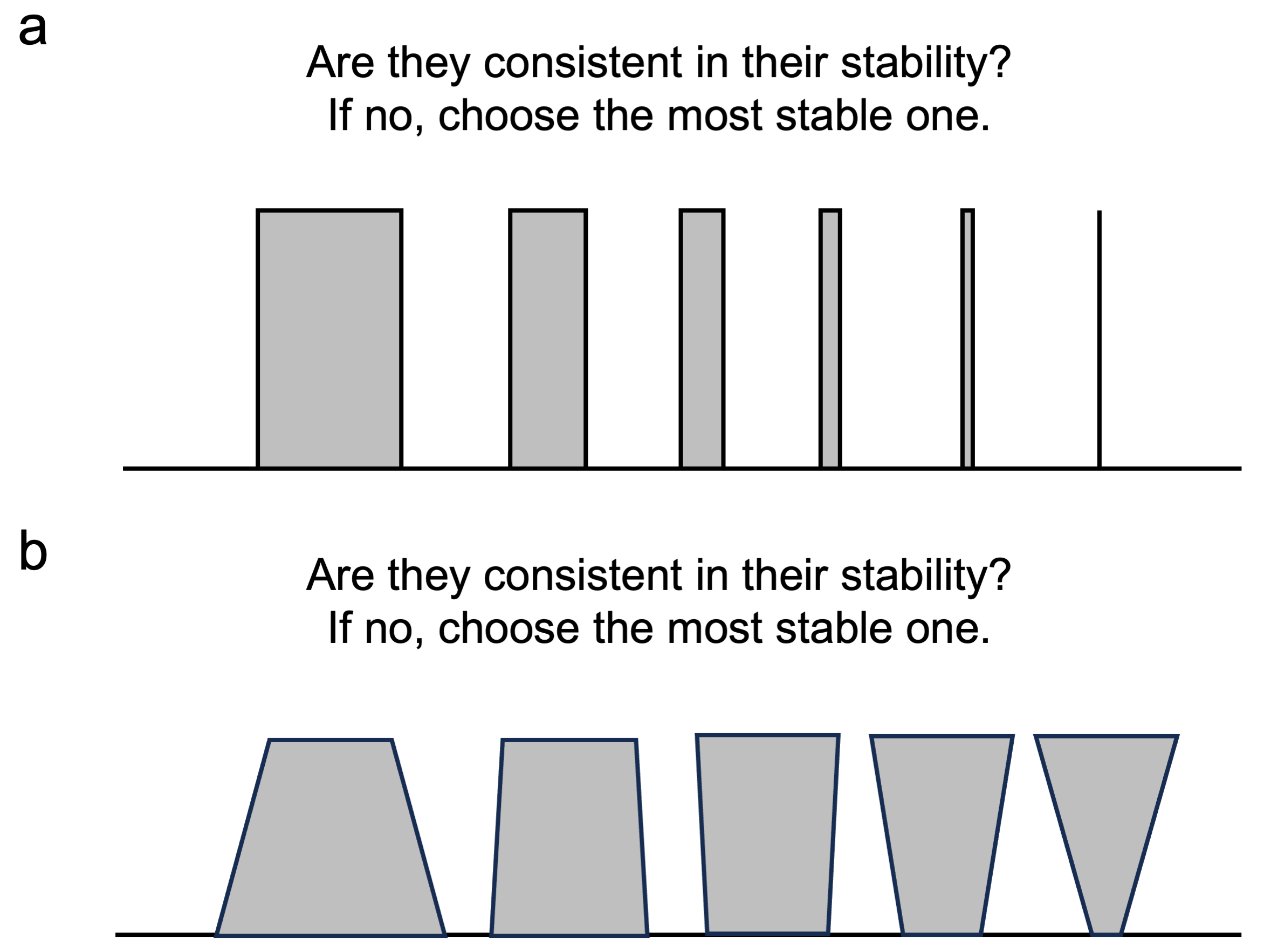

### supFigure3.png

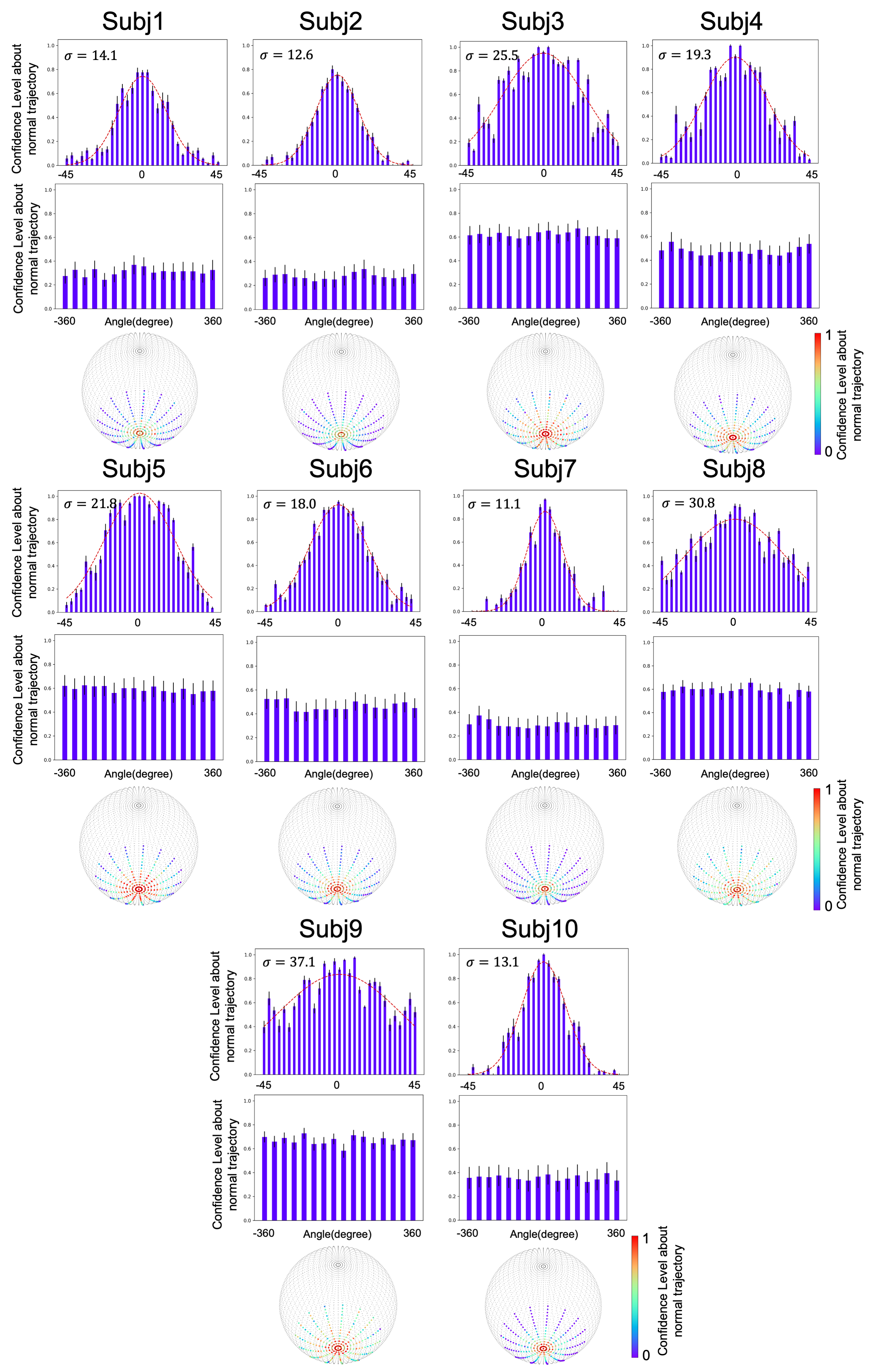

### supFigure4.png

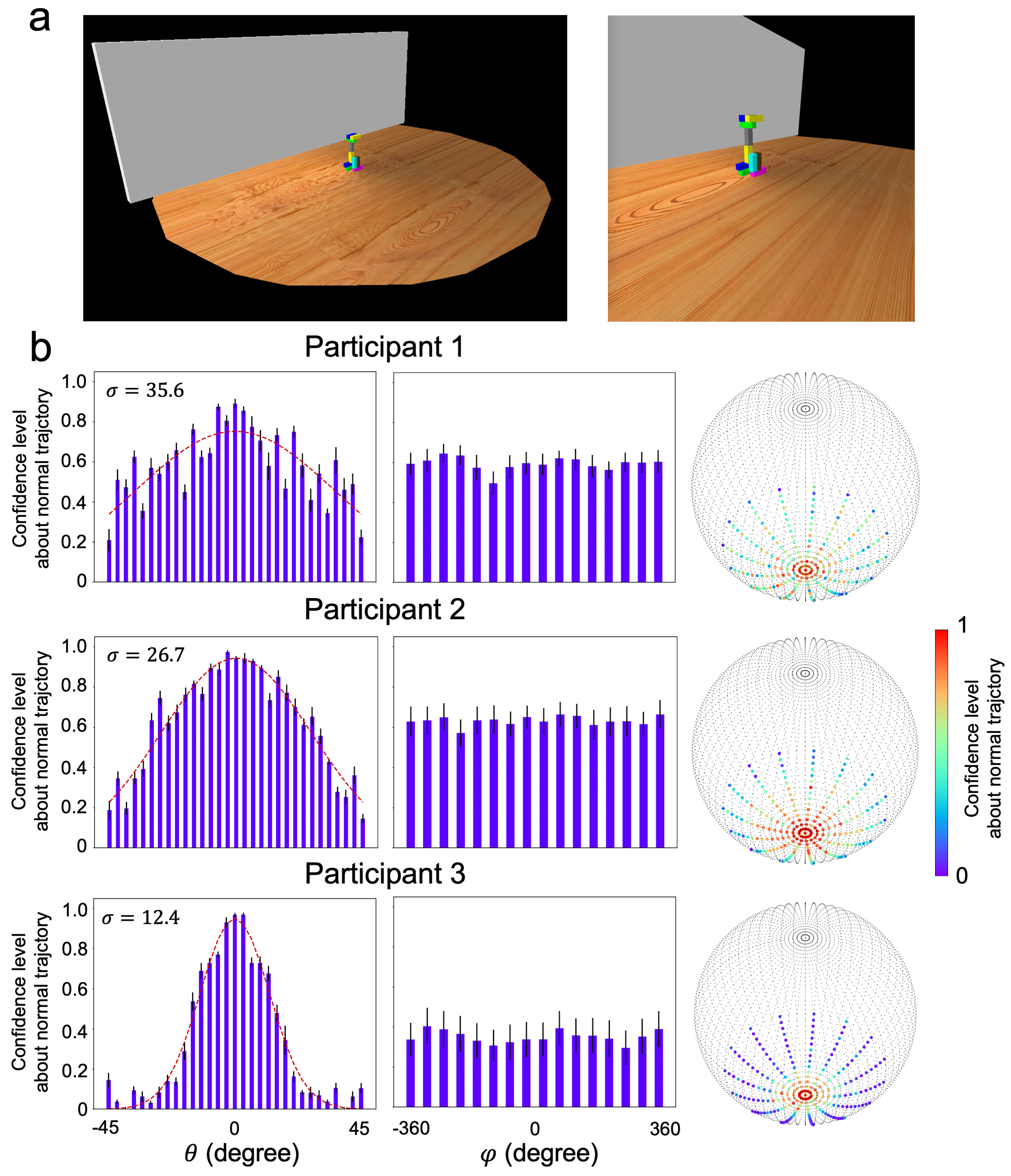

### supFigure5.png

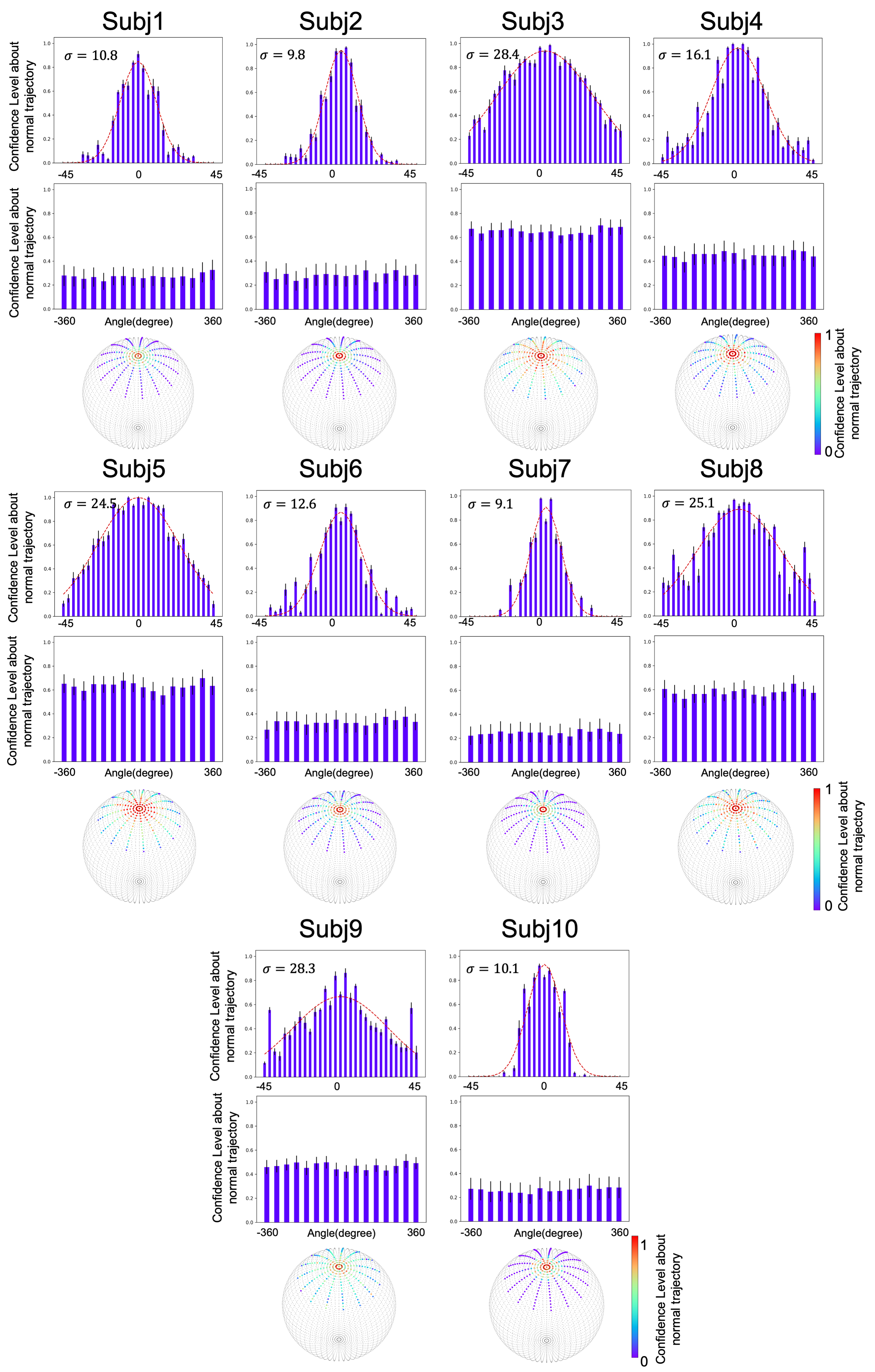

### supFigure6.png

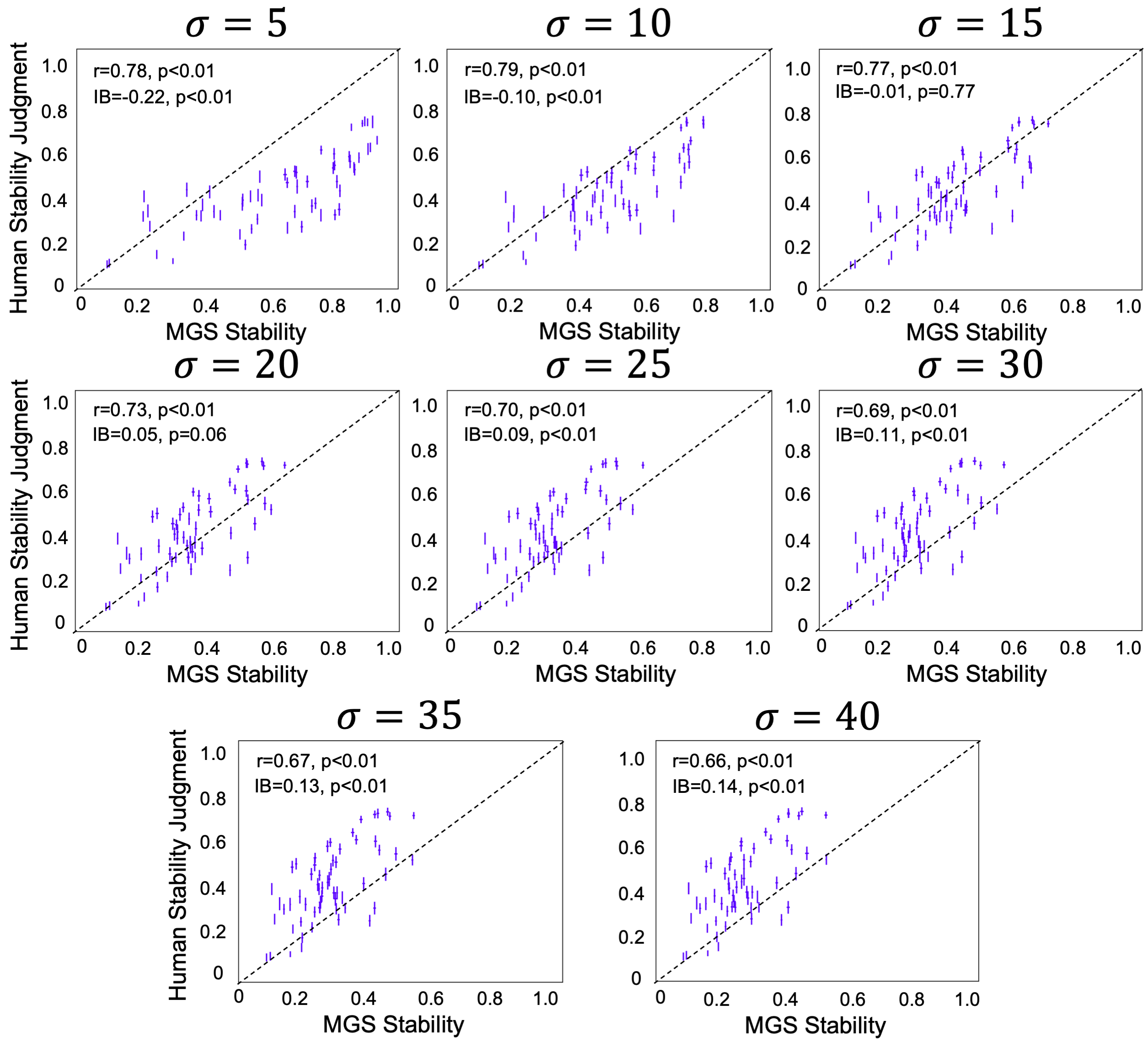

### supFigure7.png

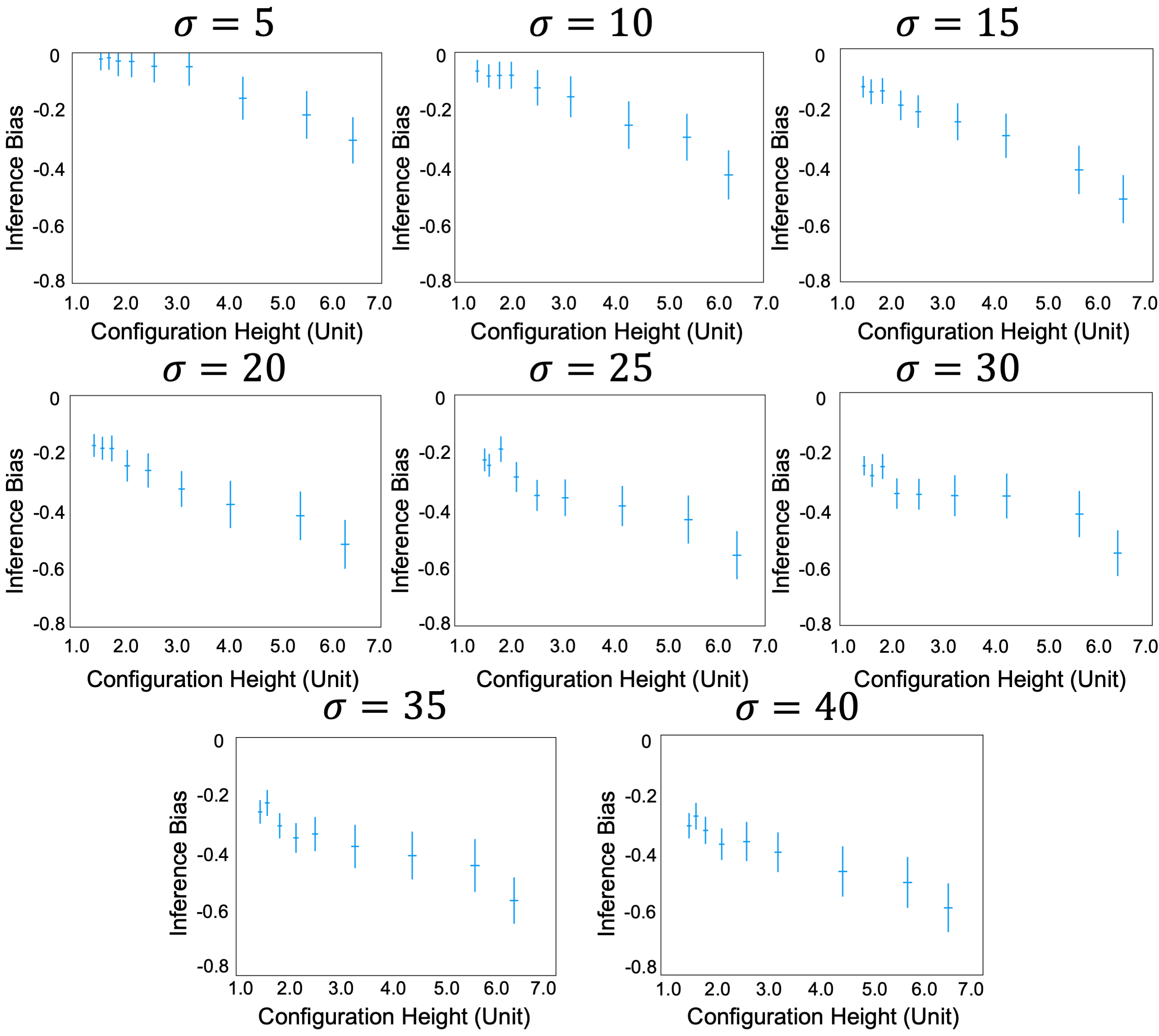

### supFigure8.png

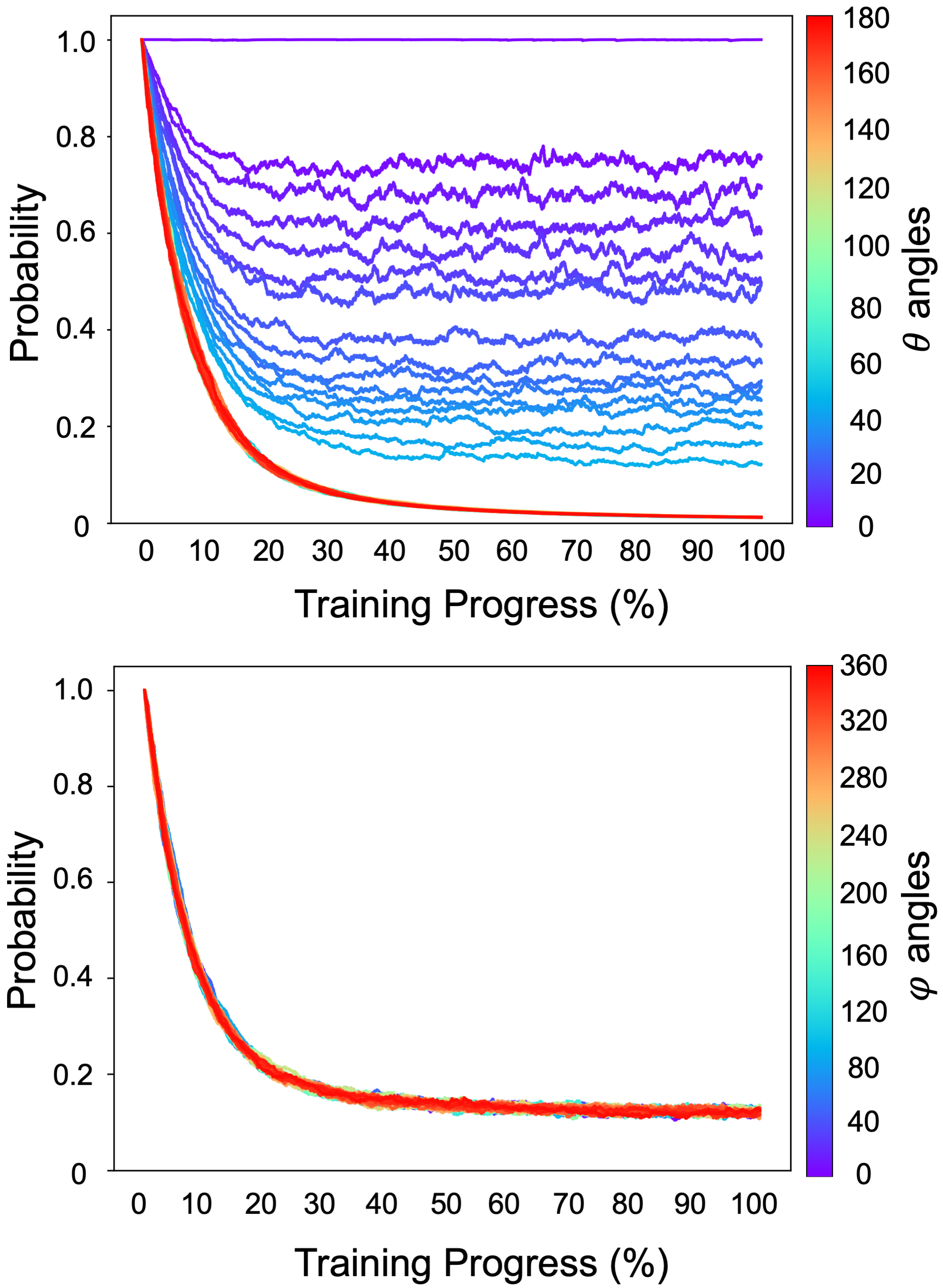

### supFigure9.png

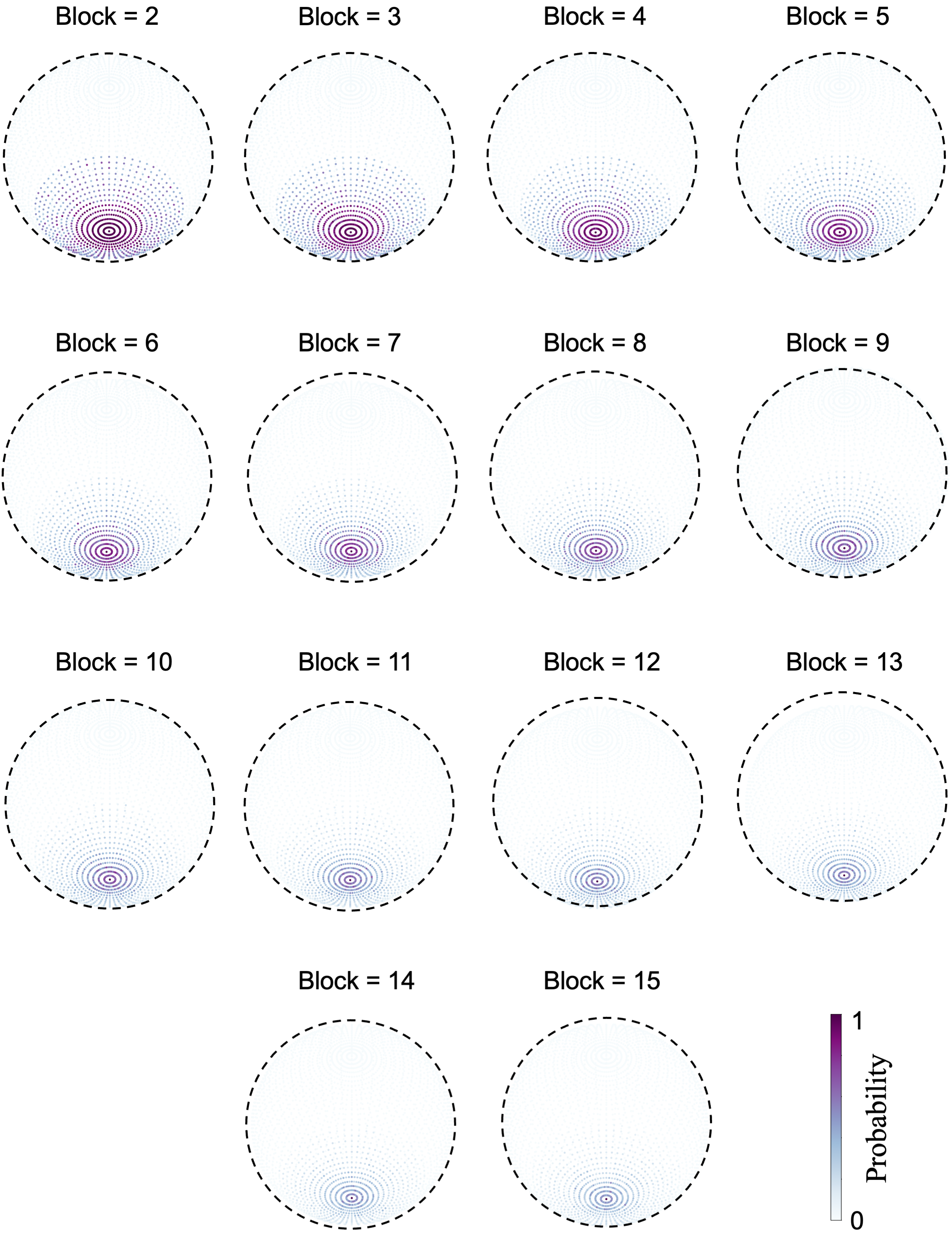

### supFigure10.png

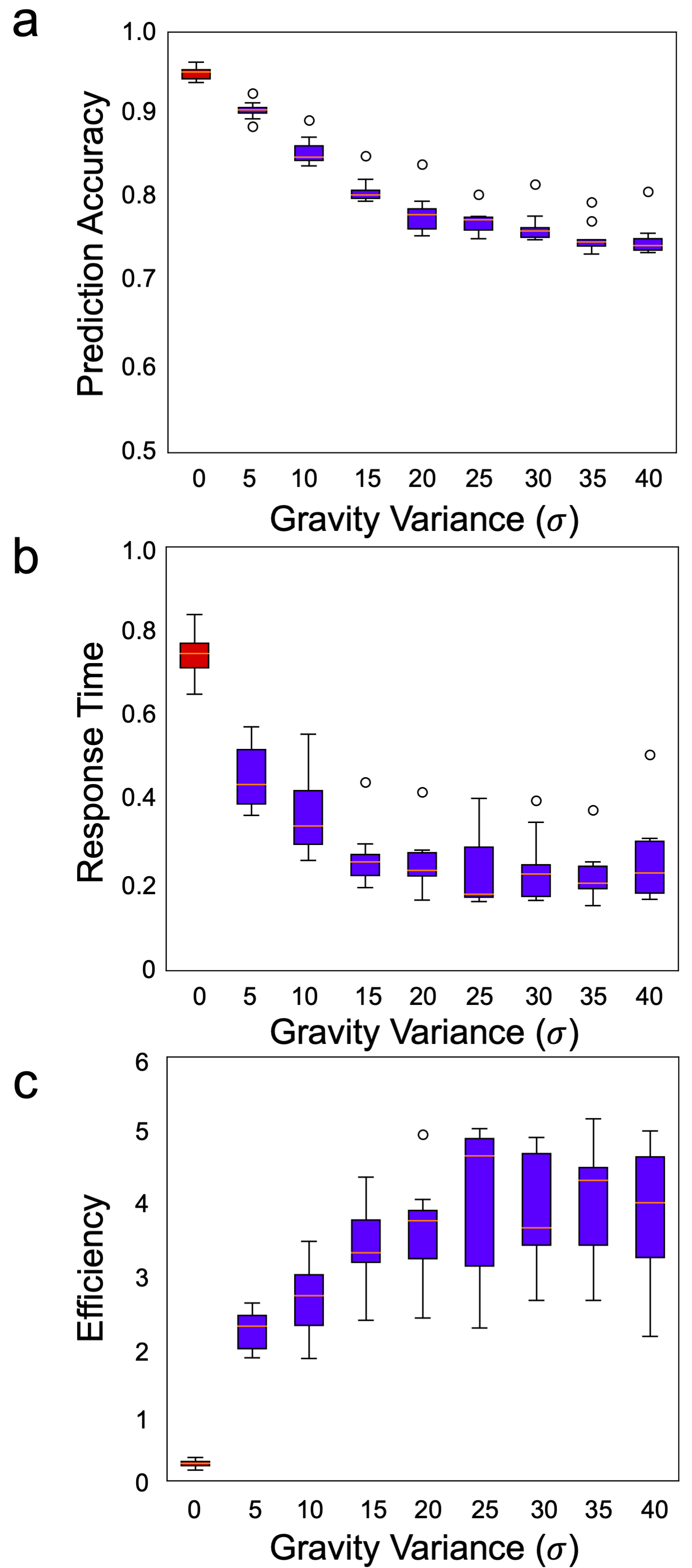

### supFigure11.png

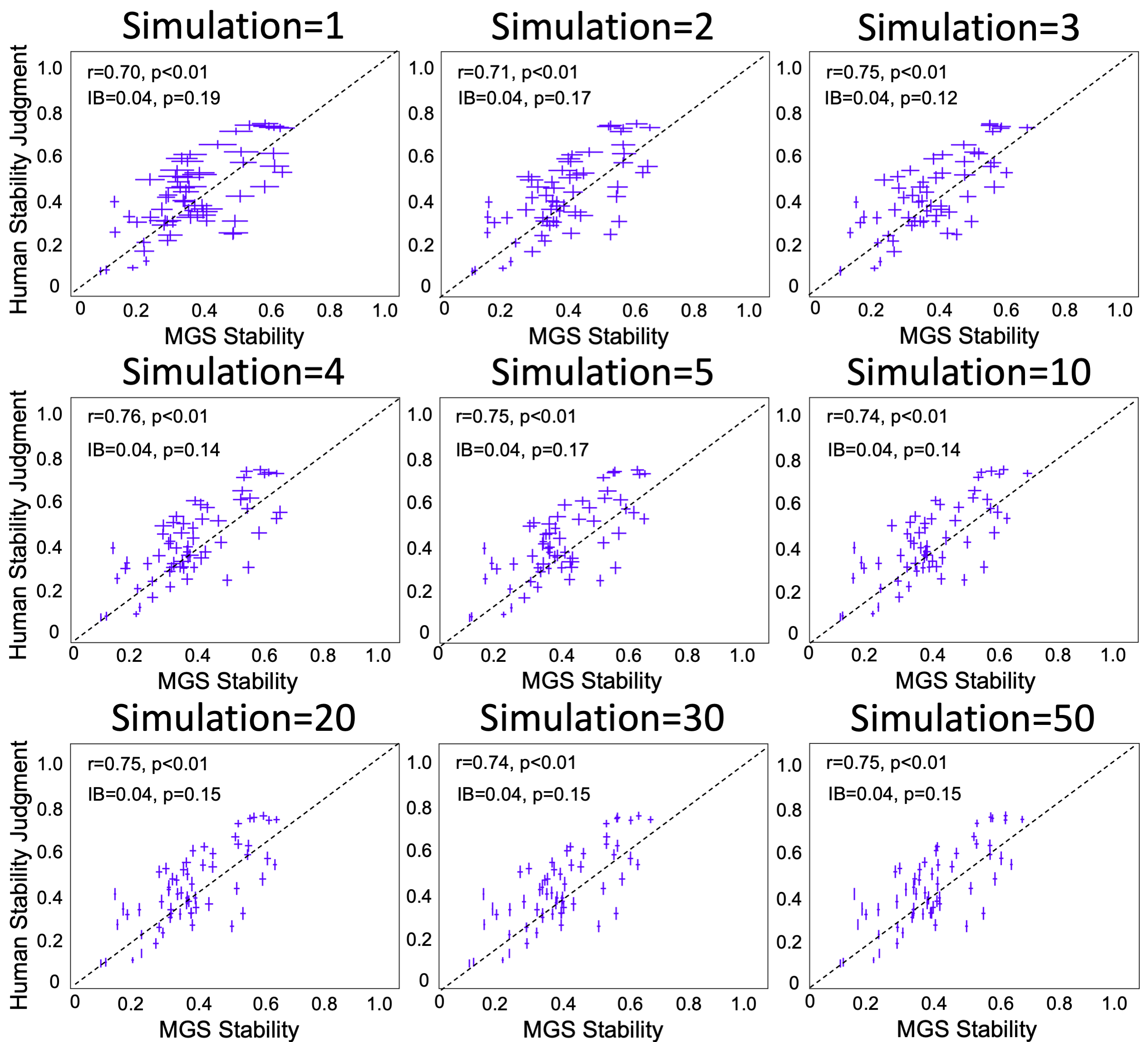
